# Estrogen-related differences in antitumor immunity and the gut microbiome contribute to sexual dimorphism of colorectal cancer

**DOI:** 10.1101/2024.01.17.576009

**Authors:** Georgia Lattanzi, Federica Perillo, Angélica Díaz-Basabe, Bruna Caridi, Chiara Amoroso, Alberto Baeri, Elisa Cirrincione, Michele Ghidini, Barbara Galassi, Elisa Cassinotti, Ludovica Baldari, Luigi Boni, Maurizio Vecchi, Flavio Caprioli, Federica Facciotti, Francesco Strati

## Abstract

Colorectal cancer (CRC) is a multifaceted disease whose development and progression varies depending on tumor location, age of patients, infiltration of immune cells within cancer lesions, and the tumor microenvironment. These pathophysiological characteristics are additionally influenced by sex-related differences. The gut microbiome plays a pivotal role in the initiation and progression of CRC, and shapes anti-tumor immune responses but how the responsiveness of the immune system to the intestinal microbiota may contribute to the sexual dimorphism of CRC is largely unknown. Here, we studied survival, tumor-infiltrating immune cell populations and tumor-associated microbiome of a cohort of n=184 male and female CRC patients and functionally tested the immune system-microbiome interactions in *in vivo* and *in vitro* models of the disease. High-dimensional single-cell flow cytometry showed that female patients are enriched by tumor-infiltrating iNKT cells but depleted by cytotoxic T lymphocytes. The enrichment of oral pathobionts and a reduction of β-glucuronidase activity are distinctive traits characterizing the gut microbiome of women affected by CRC. Functional assays using a collection of human primary iNKT cell lines demonstrated that the gut microbiota of female patients functionally impairs iNKT cell anti-tumor functions interfering with the granzyme-perforin cytotoxic pathway. These results highlight a sex-dependent functional relationship between the gut microbiome, estrogen metabolism, and the decline of cytotoxic T cell responses, contributing to the sexual dimorphism observed in CRC patients with relevant implications for precision medicine and the design of targeted therapeutic approaches addressing sex bias in cancer.

## Introduction

Colorectal cancer (CRC) is one of the most common causes of cancer morbidity both in men and in women. CRC cannot be considered merely as a type of disease; its pathogenesis depends, for example, on the anatomical location of the tumor, patients’ age, and is characterized by sex- and gender-related differences (1). Overall, sexual dimorphism exists at multiple levels in CRC. Women, compared to men, have a lower risk to develop CRC (1) although incidence and mortality in patients over 65 years old are higher in women than men (2-4). Moreover, women have a higher prevalence of right-sided colon cancer, a deadlier form of CRC, characterized by high level of microsatellite instability (MSI) that is associated with a 20% increased risk of death compared with cancer of the left side (2). However, estrogens are considered as a protective factor against MSI (5-7). Thus, the elevated risk of MSI-high colon cancer observed in older women may be associated to an age-related decline of estrogens. In line with this, the Women’s Health Initiative Clinical Trial reported a 40% reduction in colorectal cancer risk among postmenopausal women undergoing hormone replacement therapy (5, 6).

Immune infiltration in cancer lesions plays a crucial role in CRC pathophysiology (8). Generally, CRC can be classified also by the frequency and type of tumor-infiltrating immune cells (9). Patients with high infiltration of immune cells have a lower risk of disease recurrence (9). Nevertheless, the activation state of tumor-infiltrating immune cells and their production of effector molecules dictate the anti-vs pro-tumor immune responses and patient survival (8, 10). Sex hormones modulate the development and function of the immune system, shaping innate and adaptive immunity with clear consequences on antitumor responses. Indeed, estrogen’s influence on IFNγ expression and cytotoxic CD8^+^T cell infiltration in CRC indicate a more robust adaptive inflammatory T cell response in female than male patients (11). Conversely, elevated levels of intratumor neutrophils correlate with an unfavorable recurrence-free survival (12). Tumor-Associated Neutrophils (TAN) play a pivotal role in CRC acquiring a malignant phenotype and serving as an independent factor that contribute to the poor prognosis of patients (13). We recently demonstrated that tumor-infiltrating iNKT cells can contribute to the remodeling of the TME by recruiting TANs in the early phases of tumor development, thereby sculpting the CRC progression trajectory with negative outcomes in terms of patients’ overall survival (14). iNKT cells are a subset of tissue resident T cells with innate-like functions involved in tumor immune surveillance (15) endowed with killing capabilities towards CRC cells *in vitr*o (16). However, iNKT cell role in cancer biology is still controversial, possibly as a consequence of the negative impact of the TME on their functions. Indeed, we and others demonstrated that the tumor-associated microbiome and the TME impairs the antitumor functions of iNKT cells promoting immunosuppressive responses in myeloid cells (14, 17).

The gut microbiota is an oncogenic driver of CRC (18). Some tumor-associated bacterial species can promote carcinogenesis via the production of proinflammatory toxins and stimulating immunosuppressive responses (19). Dysbiosis gains further importance considering the sexual dimorphism of CRC as the gut microbiota composition changes between male and female subjects (20). Moreover, the gut microbiome affects the bioavailability of sex hormones (21). During phase II metabolism, estrogens are glucuronidated in the liver; upon entry into the gastrointestinal tract, they are exposed to microbiota-secreted β-glucuronidase that deconjugate the sugar moiety reactivating the parent compound. This allows unconjugated estrogens to be reabsorbed in the blood stream through the enterohepatic recirculation (22) suggesting a role for the estrobolome *i*.*e*., the bacterial gene repertoire affecting estrogen metabolism (23), in CRC pathophysiology. Here, by taking advantage of a large cohort of human CRC patients, murine models of colon carcinogenesis and *in vitro* functional assays, we report how estrogen-related differences in antitumor immunity and its functional interaction with the gut microbiome explain sex-dependent CRC pathogenesis.

## Results

### Decline of cytotoxic T cell responses correlates with adverse disease outcomes in women over 65 years old

Sexual dimorphism affects incidence and survival of CRC. Yet, a lack of understanding is present on which sex-related biological variables may specifically impact on cancer development, progression, and survival.

Thus, we analyzed a prospective cohort of n=184 patients enrolled at the Policlinico Hospital Milan between 2017 and 2023 to gain deeper insights into sex-related differences in survival, immunological landscape, and tumor-associated microbiota of CRC (**Figure 1A**). The clinical characteristics of CRC patients are detailed in **Table 1**. On overall, female patients showed a trend decrease in 5-year survival rate compared to men (p=0.062) and no difference in event-free survival (p=0.15). However, if stratified according to age, women over 65 years had lower 5-year and event-free survival rates compared to their age-matched male counterparts (p=0.05) (**Figure 1B-C**) while we did not observe differences in younger patients (< 65 years old). The unsupervised multidimensional immunophenotyping of freshly isolated tumor-infiltrating T cells by FACS showed a different distribution of T cell populations between male and female patients (**Figure 1D**). Tumor-infiltrating T cells belonging to clusters of T helper (C14 and C20) and cytotoxic T cells (C9) expressing IFNγ (C5) were less frequent in women than men (**Figure 1E**). Conversely, clusters of T helper cells expressing TNFα (C13), Ki67 and PD-1 (C24) as well as a cluster of iNKT cells positive for PD-1 were more frequent in women (**Figure 1E**). We further confirmed that age and the immune system are important variables when considering sex-bias in CRC by validating our results by manual gating analysis of FACS data. Indeed, we observed that CRC lesions from women were significantly infiltrated by iNKT and MAIT cells, another innate-like population of tissue resident lymphocytes, but depleted by conventional cytotoxic cells compared to age-matched men older than 65 years, as measured by reduced frequency of Natural Killer (NK) cells, conventional CD4^+^T-helper (Th), cytotoxic CD8^+^T lymphocytes (CTL) and iNKT cells expressing IFNγ and GzmA (**Figure 1F**). Thus, the decline of cytotoxic T cell responses in women may, in part, explain the reduced survival of female patients compared to man over 65 years old.

**Table 1:** Characteristics of the patient cohort.

|  | F (N=87) | M (N=97) | Total (N=184) | p value |
| --- | --- | --- | --- | --- |
| **\*\*Age\*\*** |  |  |  | 0.766 |
| Mean (SD) | 74.356 (13.353) | 73.773 (13.137) | 74.049 (13.206) |  |
| Range | 30.000 - 97.000 | 42.000 - 95.000 | 30.000 - 97.000 |  |
| **\*\*Age_at_Diagnosis\*\*** |  |  |  | 0.734 |
| Mean (SD) | 71.069 (13.360) | 70.412 (12.795) | 70.723 (13.033) |  |
| Range | 27.000 - 92.000 | 41.000 - 93.000 | 27.000 - 93.000 |  |
| **\*\*Location_General\*\*** |  |  |  | 0.053 |
| Right colon | 42 (48.3%) | 32 (33.0%) | 74 (40.2%) |  |
| Left colon | 44 (50.6%) | 65 (67.0%) | 109 (59.2%) |  |
| Multiple | 1 (1.1%) | 0 (0.0%) | 1 (0.5%) |  |
| **\*\*CT_RT_Neoadj\*\*** |  |  |  | 0.151 |
| NO | 83 (95.4%) | 86 (88.7%) | 169 (91.8%) |  |
| YES | 3 (3.4%) | 10 (10.3%) | 13 (7.1%) |  |
| YES (RT only) | 0 (0.0%) | 1 (1.0%) | 1 (0.5%) |  |
| YES (CT only) | 1 (1.1%) | 0 (0.0%) | 1 (0.5%) |  |
| **\*\*Stage\*\*** |  |  |  | 0.484 |
| 0 | 0 (0.0%) | 3 (3.1%) | 3 (1.6%) |  |
| I | 19 (21.8%) | 19 (19.6%) | 38 (20.7%) |  |
| II | 3 (3.4%) | 4 (4.1%) | 7 (3.8%) |  |
| IIA | 15 (17.2%) | 25 (25.8%) | 40 (21.7%) |  |
| IIB | 2 (2.3%) | 2 (2.1%) | 4 (2.2%) |  |
| IIC | 2 (2.3%) | 2 (2.1%) | 4 (2.2%) |  |
| IIIA | 3 (3.4%) | 5 (5.2%) | 8 (4.3%) |  |
| IIIB | 24 (27.6%) | 21 (21.6%) | 45 (24.5%) |  |
| IIIC | 7 (8.0%) | 2 (2.1%) | 9 (4.9%) |  |
| IVA | 9 (10.3%) | 7 (7.2%) | 16 (8.7%) |  |
| IVB | 1 (1.1%) | 3 (3.1%) | 4 (2.2%) |  |
| IVC | 2 (2.3%) | 4 (4.1%) | 6 (3.3%) |  |
| **\*\*Surgery\*\*** |  |  |  | 0.337 |
| Anterior_Resection | 18 (20.7%) | 26 (26.8%) | 44 (23.9%) |  |
| Hemicolectomy | 51 (58.6%) | 58 (59.8%) | 109 (59.2%) |  |
| **\*\*Tumor_Residual\*\*** |  |  |  | 0.48 |
| Complete_Resection | 83 (95.4%) | 95 (97.9%) | 178 (96.7%) |  |
| Macroscopic_Residual | 1 (1.1%) | 0 (0.0%) | 1 (0.5%) |  |
| Microscopic_Residual | 3 (3.4%) | 2 (2.1%) | 5 (2.7%) |  |
| **\*\*Vascular_Invasion\*\*** |  |  |  | 0.987 |
| NO | 61 (70.1%) | 69 (71.1%) | 130 (70.7%) |  |
| YES | 25 (28.7%) | 27 (27.8%) | 52 (28.3%) |  |
| NA | 1 (1.1%) | 1 (1.0%) | 2 (1.1%) |  |
| **\*\*Linfatic_Invasion\*\*** |  |  |  | 0.991 |
| NO | 84 (96.6%) | 94 (96.9%) | 178 (96.7%) |  |
| YES | 2 (2.3%) | 2 (2.1%) | 4 (2.2%) |  |
| NA | 1 (1.1%) | 1 (1.0%) | 2 (1.1%) |  |
| **\*\*Perineural_Invasion\*\*** |  |  |  | 0.53 |
| NO | 66 (75.9%) | 76 (78.4%) | 142 (77.2%) |  |
| YES | 20 (23.0%) | 18 (18.6%) | 38 (20.7%) |  |
| NA | 1 (1.1%) | 3 (3.1%) | 4 (2.2%) |  |
| **\*\*Budding\*\*** |  |  |  | 0.945 |
| NO | 25 (28.7%) | 28 (28.9%) | 53 (28.8%) |  |
| YES | 60 (69.0%) | 66 (68.0%) | 126 (68.5%) |  |
| NA | 2 (2.3%) | 3 (3.1%) | 5 (2.7%) |  |
| **\*\*Linfatic_Infiltration\*\*** |  |  |  | 0.008 |
| NO | 21 (24.1%) | 44 (45.4%) | 65 (35.3%) |  |
| YES | 65 (74.7%) | 51 (52.6%) | 116 (63.0%) |  |
| NA | 1 (1.1%) | 2 (2.1%) | 3 (1.6%) |  |
| **\*\*MMR_status\*\*** |  |  |  | 0.004 |
| Deficient | 23 (26.4%) | 10 (10.3%) | 33 (17.9%) |  |
| Proficient | 64 (73.6%) | 87 (89.7%) | 151 (82.1%) |  |
| **\*\*Progression\*\*** |  |  |  | 0.877 |
| NO | 65 (74.7%) | 75 (77.3%) | 140 (76.1%) |  |
| YES | 11 (12.6%) | 12 (12.4%) | 23 (12.5%) |  |
| NA | 11 (12.6%) | 10 (10.3%) | 21 (11.4%) |  |

**Figure 1:**
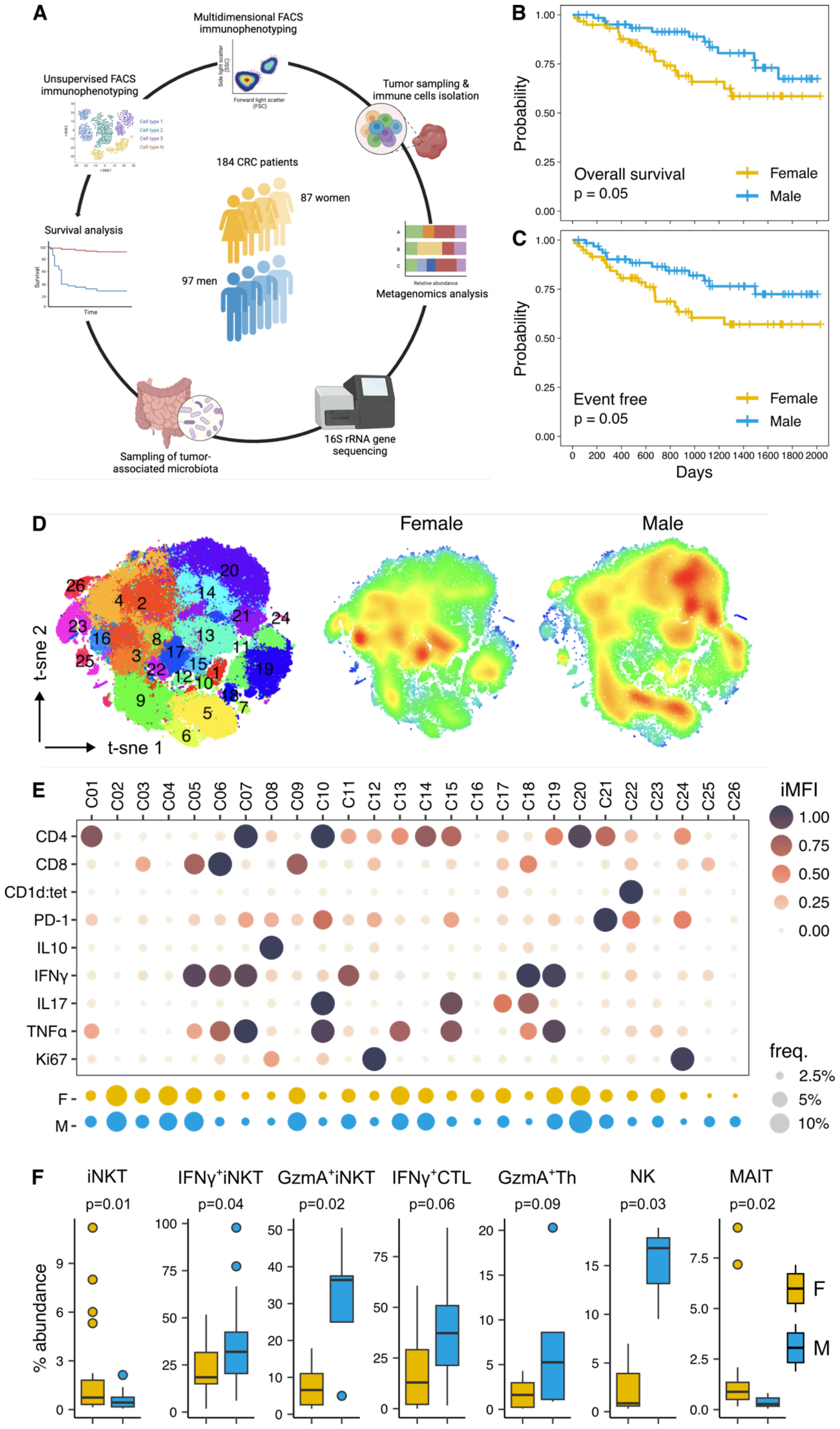
Decline of cytotoxic T cell responses in women vs men in CRC. **A)** Overview of the human study design. **B-C)** Kaplan-Meier curves for **B)** overall and **C)** event-free survival in male vs female patients with > 65 years old. **D)** t-SNE map with density plot of tumor-infiltrating CD3^+^T cells based on Phenograph clustering in male and female patients. **E)** Balloon plot of the scaled integrated MFI of Phenograph clusters generated in D. **F)** Frequency of significantly different tumor-infiltrating T cell populations in male and female patients with > 65 years old as measured by manual gating analysis of FACS data.

### The tumor-associated microbiome of female patients is rich of oncomicrobes correlating with immunosuppressive responses

The gut microbiome is an oncogenic driver of CRC (18, 19) and its composition changes according to sex (24). Thus, we characterized the tumor-associated microbiota from patients (n=35) to unravel its implication in the sexual dimorphism of CRC. Albeit, we did not find significant shifts in the overall microbial community structure between male and female patients (**Figure 2A-B**), we observed a specific sex-biased enrichment of different oncomicrobes (**Figure 2C**). Male patients were enriched by potentially genotoxic and immunogenic bacteria belonging to the genera *Bacteroides* and *Escherichia* (19). Conversely, female patients were enriched by immunosuppressive pathobionts of oral origin such as *Fusobacterium, Parvimonas, Anaerococcus* and *Alloprevotella* (25) (**Figure 2C**). Given the impact of enterohepatic recirculation on estrogen half-life and systemic levels in men and postmenopausal women (26), we hypothesized that sex-related differences in the gut microbiota composition could be associated with reduced microbial β-glucuronidase activity in women, leading to a diminished availability of systemic active estrogens. We performed a functional metagenomic prediction analysis of the microbiome revealing a significant gene content reduction of microbial β-glucuronidases (KEGG orthology K01195) in women compared to men (**Figure 2D**). Moreover, we observed the enrichment of metabolic pathways related to amino acid and lipid metabolism in women and metabolism of simple sugars like galactose, mannose, and fructose in men (**Supplementary Figure 1**). Next, we evaluated whether the tumor-associated microbiome contributed to functionally polarize specific immune responses that align with sexual dimorphism in CRC. The correlation analysis of the most abundant bacterial taxa (mean abundance > 0.25%) with the frequency of tumor-infiltrating immune cells revealed distinct clusters of bacteria showing positive or negative correlations with antigen presenting cells (APC) expressing MHCII and CD1d (**Figure 2E**). Interestingly, these bacterial clusters also exhibited corresponding positive or negative correlations with type 17 and type 2 immune responses and T cells expressing the immune checkpoint CTLA-4, aligning inversely with the correlations observed for APCs (**Figure 2E**).

**Figure 2:**
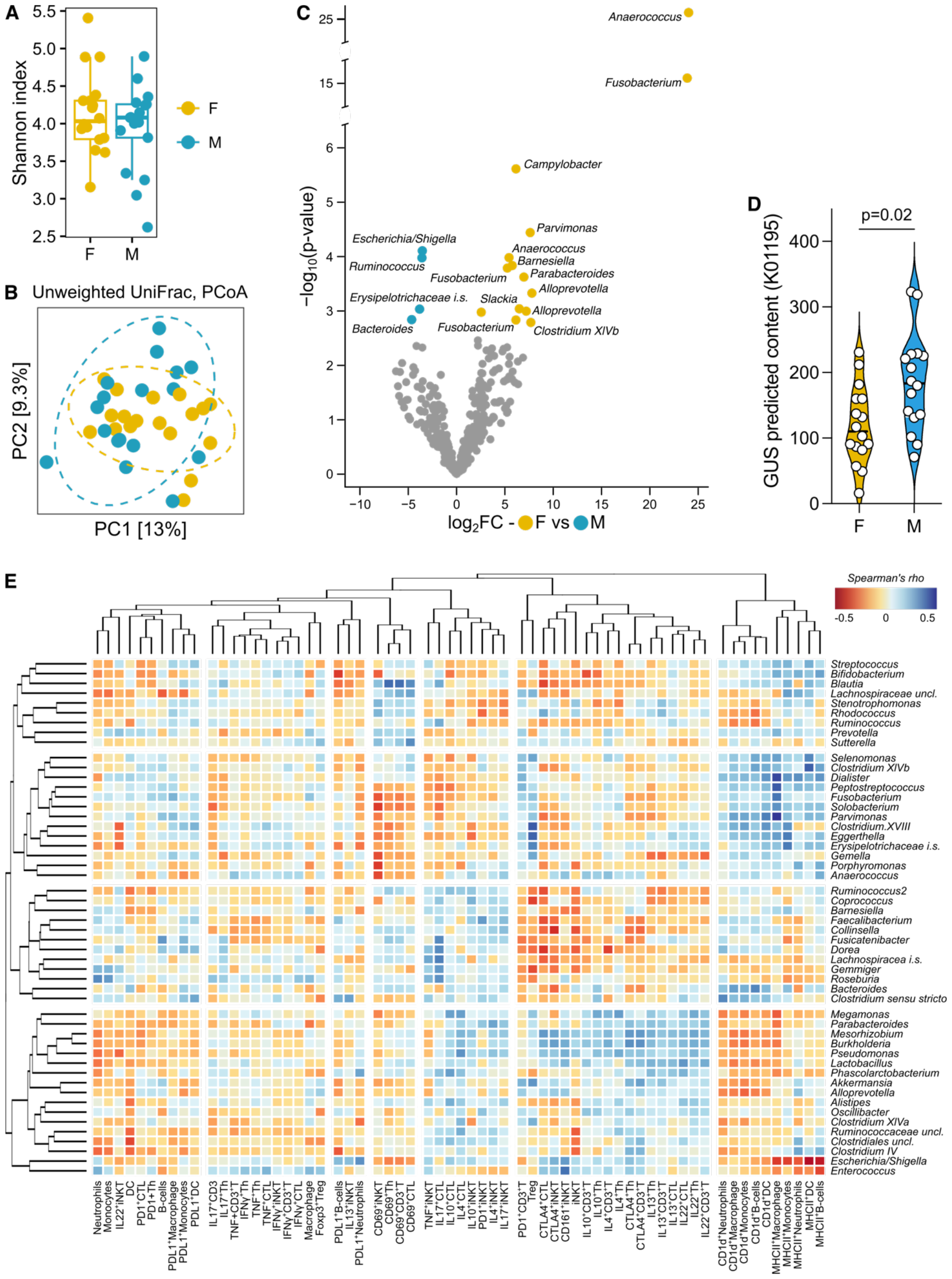
Tumor-associated immune-microbiota interactions in CRC patients. **A)** Microbiota α-diversity as measured by Shannon index in female and male CRC patients. **B)** PCoA of microbial β-diversity as measured by unweighted UniFrac distance. **C)** Volcano plot showing the significantly enriched bacterial amplicon sequence variants (ASVs) (p<0.05) by the DEseq2 analysis. The names of the significantly enriched bacterial ASVs classified up to the genus level are reported. **D)** Predicted metagenomic gene content of β-glucuronidase (GUS) inferred by Tax4Fun2 analysis. **E)** Heatmap of Spearman’s rho correlations between the relative abundance of the most abundant (mean relative abundance > 0.25%) tumor-associated bacterial taxa with the frequency of tumor-infiltrating immune cells analyzed by FACS in male and female CRC patients. CTL, CD8^+^ cytotoxic T lymphocytes; Th, CD4^+^ T helper cells; DC, dendritic cells.

A cluster of oral pathogens, including *Fusobacterium, Porphyromonas, Peptostreptococcus*, and *Parvimonas*, exhibited a positive correlation with the frequency of APCs, T cells expressing PD-1, and T-regulatory (Treg) cells. Conversely, this bacterial cluster showed a negative correlation with activated/tissue-resident CD69^+^T cells (**Figure 2E**). Altogether, these data suggests that sex-based differences in the tumor-associated microbiome and its metabolism may influence diverse immune responses potentially contributing to sexual dimorphism of CRC.

### Tumor progression and estrogen-related differences in immunity and microbiome composition promote sexual dimorphism of CRC in the AOM/DSS mouse model

To better understand sex-biases influencing the developmental pathway of CRC, we compared tumor burden and immunophenotypes between female and male animals in the AOM/DSS colon cancer model at different time-point of tumorigenesis (**Figure 3A**). We observed a worse endoscopic score and a higher number of tumors in males compared to females at an early timepoint (T1) following the initiation of tumorigenesis (**Figure 3B-D**). However, the disparity in tumor burden significantly diminished at a later timepoint (T2) (**Figure 3C-D**), suggesting that aging in female mice may influence CRC progression more than in males. Immunophenotyping of tumor-infiltrating cells revealed a reduction of conventional CD4^+^ helper and CD8^+^ cytotoxic T cells at T2 compared to T1 in female mice (**Figure 3E-G**). In contrast, CD8^+^T cells increased over time in male mice, suggesting a progressive decline in antitumor immunity in females, as observed in CRC patients. Furthermore, we observed a consistently higher frequency of iNKT cells in tumor-bearing female mice at both T1 and T2 (**Figure 3H-I**), mirroring the observations in human patients. As alterations in gut microbiome composition can impact disease progression, we performed a metagenomic analysis of female tumor-bearing C57BL/6 mice at both timepoints from the induction of CRC. The gut microbiome of AOM/DSS mice exhibited a reduction of microbial α-diversity compared to controls (p=0.132; AOM vs CTRL, T1; p=0.027; AOM vs CTRL, T2) (**Figure 3J**) and significant changes of microbiome composition as the disease progressed, as measured by β-diversity (unweighted UniFrac, R^2^=0.19, F-stat=2.11, p=0.021; AOM/DSS, T1 vs T2) (**Figure 3K**) and DESeq2 analysis (**Figure 3L**).

**Figure 3:**
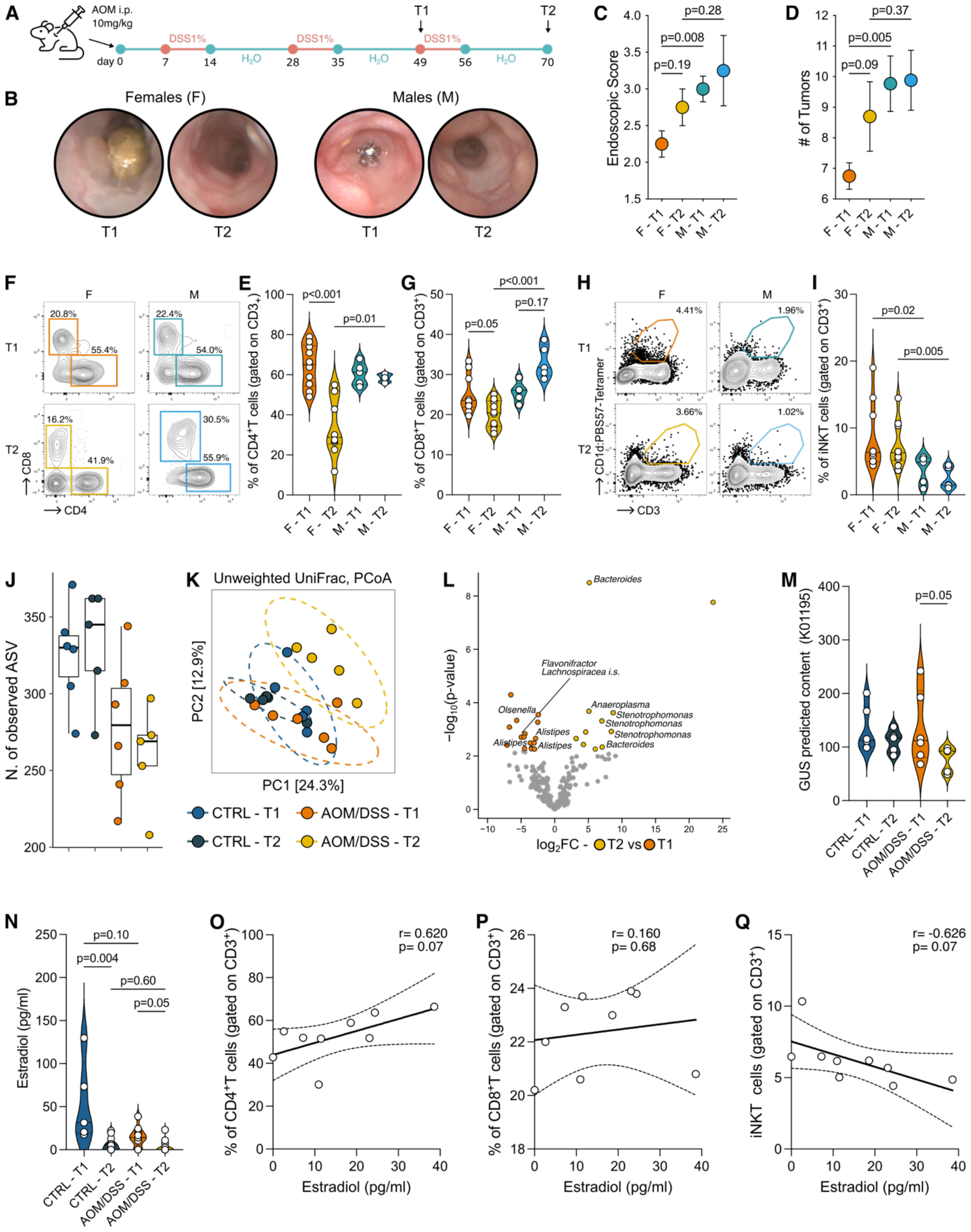
Age-related decline of estrogen levels and antitumor immunity correlates with reduced protection from colorectal tumorigenesis in female tumor-bearing mice. **A)** Schematic representation for the AOM/DSS experimental plan. **B-C)** Tumor endoscopic score and representative endoscopic pictures; **D)** number of tumors from male (M) and female (F) AOM/DSS treated C57BL/6 animals at early (T1) and late (T2) timepoints from the induction of tumorigenesis. **E-G)** Frequency of tumor-infiltrating **F)** CD4^+^ and **G)** CD8^+^ T cells in the AOM/DSS model with representative plots. **H-I)** Frequency of tumor-infiltrating iNKT cells with representative plots. **J)** Observed number of ASVs in female AOM/DSS treated C57BL/6 animals at early (T1) and late (T2) timepoints from the induction of tumorigenesis and age-matched control mice (CTRL). **K)** PCoA of microbial β-diversity as measured by Unweighted UniFrac distance. **L)** Volcano plot showing the significantly enriched bacterial amplicon sequence variants (ASVs) (p<0.05) by the DEseq2 analysis. The names of the significantly enriched bacterial ASVs classified up to the genus level are reported. **M)** Predicted metagenomic gene content of β-glucuronidase (GUS) inferred by Tax4fun2 analysis. **N)** measurement of serum estrogen by quantitative ELISA. **O-Q)** Correlation of serum estrogen with tumor-infiltrating **O)** CD4^+^T, **P)** CD8^+^T and **Q)** iNKT cells. Data points, n=5-10 from two pooled independent experiments representative of three.

The microbial community structure of tumor-bearing female mice at T1 demonstrated a greater similarity with tumor-free control animals (CTRL) than AOM/DSS mice at T2 (**Figure 3K**). Notably, tumor-bearing animals showed a greater reduction in β-glucuronidase activity (KEGG orthology K01195) at T2 compared to T1 and CTRL animals, as measured by the analysis of predicted microbiome functions (**Figure 3M**). The analysis of serum estradiol showed that its level decrease over time and is significantly lower in tumor-bearing mice compared to age-matched controls (**Figure 3N**). In tumor-bearing mice, estradiol appeared to modulate T cell infiltration, as shown by the positive correlation with CD4^+^T cells (**Figure 3O**) and the negative correlation with iNKT cells (**Figure 3Q**). Taken together, these findings suggest an intertwined relationship among the gut microbiome, estrogen metabolism, and antitumor immune responses.

### The gut microbiome affects iNKT cell cytotoxic functions in a disease and sex dependent manner

iNKT cells are potentially cytotoxic cells and we found them significantly enriched in tumors of female patients (**Figure 1F**). Nevertheless, iNKT cell infiltration correlates with protumor responses and negative outcomes in CRC because of the detrimental effects exerted by *Fusobacterium nucleatum* (14). Thus, we performed a series of experiments involving the priming of intestinal and circulating human iNKT cell lines (16, 27) with monocyte-derived dendritic cells (moDC) loaded with the fecal microbiota from CRC patients (**Figure 4A**) to test the hypothesis that iNKT antitumor functions could be impaired by the tumor-associated microbiome in a sex-dependent fashion. We observed that the microbiome from female patients significantly reduced the iNKT cell-mediated killing of CRC cells compared to both male patients and healthy controls (**Figure 4B**). We observed also a reduced release of granzyme (**Figure 4C**) and perforin (**Figure 4D**) upon stimulation with the microbiome from female patients in agreement with our previous demonstration that iNKT cells depend on the perforin–granzyme pathway for proper elimination of colon cancer cells (16). Altogether these data suggest that iNKT cells may play a role in CRC pathophysiology in women.

**Figure 4:**
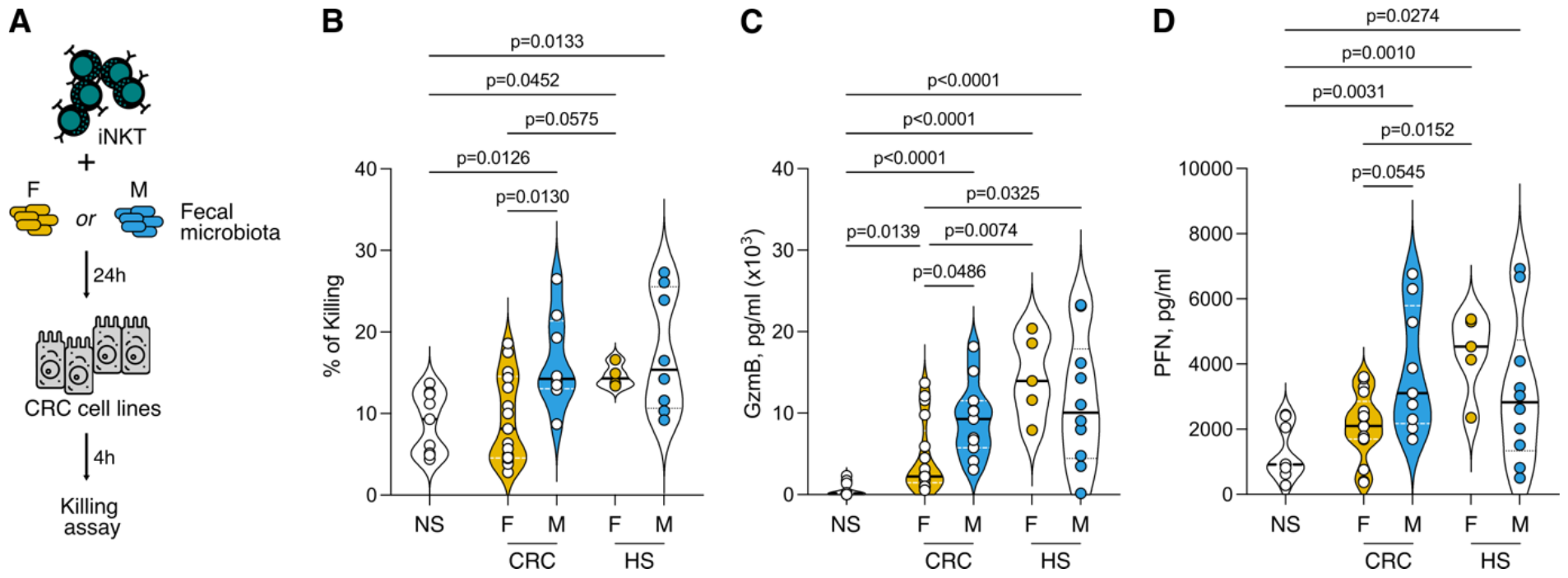
The fecal microbiota of female CRC patients impairs iNKT cell cytotoxicity. **A)** Schematic representation of the experimental plan. **B)** Percentage of killed tumor cells by iNKT cells activated by the fecal microbiota of female (F) and male (M) CRC patients and healthy subjects (HS). **C)** Granzyme B (GzmB) and **D)** perforin (PFN) concentration in the culture supernatant of iNKT cells activated by the fecal microbiota of female (F) and male (M) CRC patients and healthy subjects (HS). NS, non-stimulated. Data are representative of at least three independent experiments.

## Discussion

The mortality rate for CRC is higher in men than in women in the general population (1), creating a misconception that CRC is not a primary health concern for women. Contrary to this belief, CRC is the second cause for cancer death in women and, among individuals aged 65 and older, women exhibit lower survival rates than men, as observed in our study cohort (**Figure 1B-C**) and in previous studies (2-4). This highlights the potential influence of sex hormones through various mechanisms. Despite the established carcinogenic role of sex hormones in many tissues, they exert protective effects in the colon (28). This could be explained by the role of sex hormones in shaping immune responses, and possibly as a consequence of their interaction with the gut microbiome. Indeed, by deconjugating estrogens, the gut microbiome can affect estrogen bioavailability locally and systemically (23). Dysbiosis *i*.*e*., alteration in the microbiome composition, can disrupt this process, potentially reducing estrogen levels and diminishing their protective role in colorectal carcinogenesis. In this study, we demonstrated sex-specific alterations in the tumor-associated microbiome in CRC patients, with a notable reduction in microbial β-glucuronidase functions in women (**Figure 2C-D**). This reduction has important functional consequences as estrogens regulate the expression of cytotoxic effector molecules, such as IFNγ and granzyme A, in CD8^+^T (11) and iNKT cells (29). The decrease expression of these effector molecules can result in a decline of intratumor cytotoxic T cell responses, as shown by the immunophenotyping of older women compared to age-matched men (**Figure 1F**). Additionally, in the AOM/DSS animal model of CRC, a decline in β-glucuronidase activity and circulating estrogens was observed as the disease progressed in female mice, accompanied by alterations in the composition of the gut microbiota (**Figure 3J-N**). Consistent with our findings, a recent study using this model showed that intestinal deletion of the estrogen receptor ERβ exacerbates the severity of CRC and affects gut microbiome composition (30). How sensing of estrogens may influence the gut microbiome has not been demonstrated in this study but it is plausible to speculate that the immune system may be involved in this process. Indeed, there is a bidirectional crosstalk between the gut microbiota and the immune system that is important in health and disease (31). When correlating the gut microbiome with tumor-infiltrating immune cell populations, we observed that women, compared to men, are enriched by oncomicrobes aligning with an immunosuppressive response (**Figure 2E**). Among the microbial taxa enriched in female patients, it is worth to mention *Fusobacterium*. The periodontal pathobiont *Fusobacterium nucleatum* is considered a risk factor for CRC. It has the potential to stimulate the proliferation of cancer cells, reshape the immune microenvironment, and contribute to metastasis and chemoresistance during the initiation and progression of CRC (32). Moreover, *F. nucleatum* skews iNKT cell functions towards protumor immunosuppressive responses (14) rather than promoting antitumor cytotoxic immunity in CRC (16). Notably, iNKT cells and *Fusobacterium* were enriched in female patients and when stimulated with a microbiome derived from these patients, iNKT cells lost their cytotoxic properties (**Figure 4B**).

In conclusion, our study emphasizes how the complexity of CRC is influenced by sex-biased factors such as hormones, immune cell infiltration, and the microbiome. By uncovering sex-specific mechanisms involving the interaction between the gut microbiome and cytotoxic T cell responses, our findings pave the way to design targeted therapeutic approaches in precision medicine that address sex bias in CRC with the aim to advance the efficacy of cancer treatments and improve patient outcomes.

## Material and Methods

### Human Samples

Tumor samples were collected with informed consent from patients (n = 184) diagnosed with colorectal adenocarcinoma between October 2017 and April 2023 undergoing surgical resection at IRCCS Policlinico Ospedale Maggiore, Milan, Italy, as approved by the Institutional Review Board (Milan, Area B) with permission number 566_2015.

### Isolation of tumor-infiltrating cells

Tumor samples were taken transversally to collect both marginal and core tumor zone. Human lamina propria mononuclear cells (LPMCs) were isolated as previously described (33). Briefly, the dissected intestinal mucosa was freed of mucus and epithelial cells in sequential steps with DTT (0.1 mmol/l) and EDTA (1 mmol/l) (Sigma-Aldrich) and then digested with collagenase D (400 U/ml) (Worthington Biochemical Corporation) for 5 h at 37°C in agitation. LPMCs were then separated with a Percoll gradient.

### Mice

C57BL/6 mice were housed and bred at the European Institute of Oncology (IEO) animal facility (Milan, Italy) in SPF conditions. Sample size was chosen based on previous experience. No sample exclusion criteria were applied. No method of randomization was used during group allocation, and investigators were not blinded. Age-matched male and female mice were used for experiments. Animal experimentation was approved by the Italian Ministry of Health (Auth. 10/21 and Auth. 1217/20) and by the animal welfare committee (OPBA) of the European Institute of Oncology (IEO), Italy.

### Animal experiments

7-8 weeks old mice were injected intraperitoneally with azoxymethane (AOM, Merck) dissolved in isotonic saline solution at a concentration of 10 mg/kg body weight. After 7 days, mice were given 1% (w/v) dextran sodium sulfate (DSS MW 40 kD; TdB Consultancy) in their drinking water for 7 days followed by 14 days of recovery. The cycles were repeated for a total of 2/3 DSS cycles, and mice sacrificed at day 49 and 70.

### Murine colonoscopy

Colonoscopy was performed using the Coloview system (TP100 Karl Storz, Germany). Tumor endoscopic score has been quantified as previously described (34). During the endoscopic procedure mice were anesthetized with 3% isoflurane.

### Murine cells isolation

Single-cell suspensions were prepared from the colon of C57BL/6 mice as previously described (35). Briefly, cells were isolated via incubation with 5 mM EDTA at 37°C for 30 min, followed by mechanical disruption with GentleMACS (Miltenyi Biotec). After filtration with 100-μm and 70-μm nylon strainers (BD), the LPMC were counted and stained for immunophenotyping.

### ELISA assay

Detection of murine estrogens in serum was performed using the Mouse Estrogen ELISA Kit (Abcam) according to manufacturers’ instructions.

### iNKT cell cytotoxicity assay

The human iNKT cell lines used in this study were generated as previously described (27). All cells were maintained in a humidified incubator with 95% air, 5% CO_2_ at 37ºC. iNKT cell cytotoxicity toward the human CRC cell line RKO (American Type Culture Collection, ATCC) was performed as previously described (16).

### iNKT cell conditioning by human gut microbiota

Monocyte derived dendritic cells (moDCs) were pulsed with heat-inactivated gut microbiota of female and male CRC patients as well as healthy controls and co-cultured with iNKT cells (2 × 10^5^ cells) in a 2:1:10 iNKT:moDC:microbiota ratio in RPMI-1640 supplemented with 10%FBS, Pen/Strep. After 24 h, iNKT cell activation status was estimated by extracellular or intracellular staining. Fecal samples were resuspended 1:10 (w/v) in PBS and filtered through a 0.75 μm filter to remove large debris; microbiota cell density was quantified by qPCR (36), adjusted to 2 × 10^7^ CFU·mL^−1^ and heat-killed before being stored at -80°C until use in downstream experimentation.

### Flow Cytometry

Cells were stained with labeled antibodies diluted in PBS with 1% heat-inactivated fetal bovine serum (FBS) for 20 min at 4°C. iNKT cells were stained and identified using human or mouse CD1d:PBS57 Tetramer (NIH Tetramer core facility) diluted in PBS with 1% heat-inactivated FBS for 30 min at 4°C. For intracellular cytokine labelling cells were incubated for 3 h at 37°C in RPMI-1640+10% FBS with PMA (50ng/ml, Merck), Ionomycin (1μg/ml, Merck) and Brefeldin A (10 μg/ml, Merck). Before intracellular staining, cells were fixed and permeabilized using Cytofix/Cytoperm (BD). Samples were analyzed with a FACSCelesta flow cytometer (BD Biosciences, Franklin Lakes NJ, USA) and the FlowJo software (Version 10.8, TreeStar, Ashland, OR, USA). For the multidimensional analysis using t-SNE visualization and Phenograph clustering (37), FCS files were quality checked for live, singlets and antibody agglomerates and normalized to avoid batch effects. Data were cleaned for antibodies aggregates by checking each parameter in a bimodal plot. Gate on singlets, on viable lymphocytes and subsequently on CD3^+^cells were applied. CD3^+^ populations were down-sampled to 5000 events per sample using the DownSample plugin (Version 3.3.1) of FlowJo to create uniform population sizes. Down-sampled populations were exported as FCS files with applied compensation correction. Files were then uploaded to RStudio environment (Version 3.5.3) using the flowCore package (Version 1.38.2). Data were transformed using logicleTransform() function present in the flowCore package. To equalize the contribution of each marker they were interrogated for their density distribution using the densityplot() function of the flowViz package (Version 1.36.2). Each marker was normalized using the Per-channel normalization based on landmark registration using the gaussNorm() function present in the package flowStats (Version 3.30.0). Peak.density, peak.distance and number of peaks were chosen according to each marker expression. Normalized files were analyzed using the cytofkit package through the cytofkit_GUI interface. For data visualization we used the t-Distributed Stochastic Neighbor Embedding (t-SNE) method, while for clustering we used the Phenograph algorithm. t-SNE plots were visualized on the cytofkitShinyAPP with the following parameters: perplexity=50, iterations=1000, seed=42, k=50. FCS for each cluster were generated and re-imported in FlowJo to be manually analyzed for the determination of the integrated MFI. The iMFI of different markers was scaled from 0 to 1 and used to identify Phenograph clusters (37).

### 16S rRNA gene sequencing and data analysis

Intestinal mucus scraped from human tumor lesions and murine feces were stored at -80°C until DNA extraction. DNA extraction, 16S rRNA gene amplification, purification, library preparation and pair-end sequencing on the Illumina MiSeq platform were performed as previously described (35). Reads were pre-processed using the MICCA pipeline (v.1.7.0) (http://www.micca.org) (38). Forward and reverse primers trimming and quality filtering were performed using micca trim and micca filter, respectively. Filtered sequences were denoised using the UNOISE (39) algorithm implemented in micca otu to determine true biological sequences at the single nucleotide resolution by generating amplicon sequence variants (ASV). Bacterial ASVs were taxonomically classified using micca classify and the Ribosomal Database Project (RDP) Classifier v2.13 (40). Multiple sequence alignment (MSA) of 16S rRNA gene sequences was performed using the Nearest Alignment Space Termination (NAST) algorithm (41) implemented in micca msa with the template alignment clustered at 97% similarity of the Greengenes database (42) (release 13_08). Phylogenetic trees were inferred using micca tree (43). Sampling heterogeneity was reduced rarefying samples at the depth of the less abundant sample using micca tablerare. Alpha (within-sample richness) and beta-diversity (between-sample dissimilarity) estimates were computed using the phyloseq R package (44). Permutational multivariate analysis of variance (PERMANOVA) test was performed using the adonis2 function in the vegan R package with 999 permutations. ASVs differential abundance testing was carried out using the R package DESeq2 (45) using the non-rarefied data (46). Spearman’s correlations were tested using the psych R package (47). Prediction of functional metagenomic content was inferred by using Tax4Fun2 (48) and the reference curated databases from Kyoto Encyclopedia of Genes and Genomes (KEGG) (49) implemented within the MicrobiomeAnalyst pipeline (50). Metabolic pathway maps were visualized using iPATH 3 (51).

### Statistical analysis

Statistical tests were conducted using Prism (Version 9.5.1, GraphPad) and R (version 4.3.1). Spearman’s correlation coefficient was used for the analysis of correlations. Kaplan-Meier analysis was carried out using the R packages survival (version 3-2-11) and survminer (version 0.4.9).

## Authors contribution

FS conceived the study. FF, FS, and GL designed the experiments. GL, ADB, FP, CA, BC, EC and AB performed the experiments. FF and FS supervised the experiments. FC, MV and LB contributed with reagents and resources. GL performed multidimensional FACS data analysis. FS performed 16S rRNA sequencing and statistical data analyses. FS wrote the manuscript. All authors reviewed and critically edited the manuscript. All authors contributed to the article and approved the submitted version.

## Acknowledgments

We thank the IEO Animal Facility for the excellent animal husbandry and the NIH Tetramer Facility for providing human and murine CD1d:PBS57 tetramers. We thank Maria Rita Giuffrè and Luca Iachini for the assistance with the experiments. Panels 3A and 5A were created using icons from the Noun Project (https://thenounproject.com/). Panel 1A was created with BioRender (https://biorender.com/).

---

**Supplementary Figure 1:**
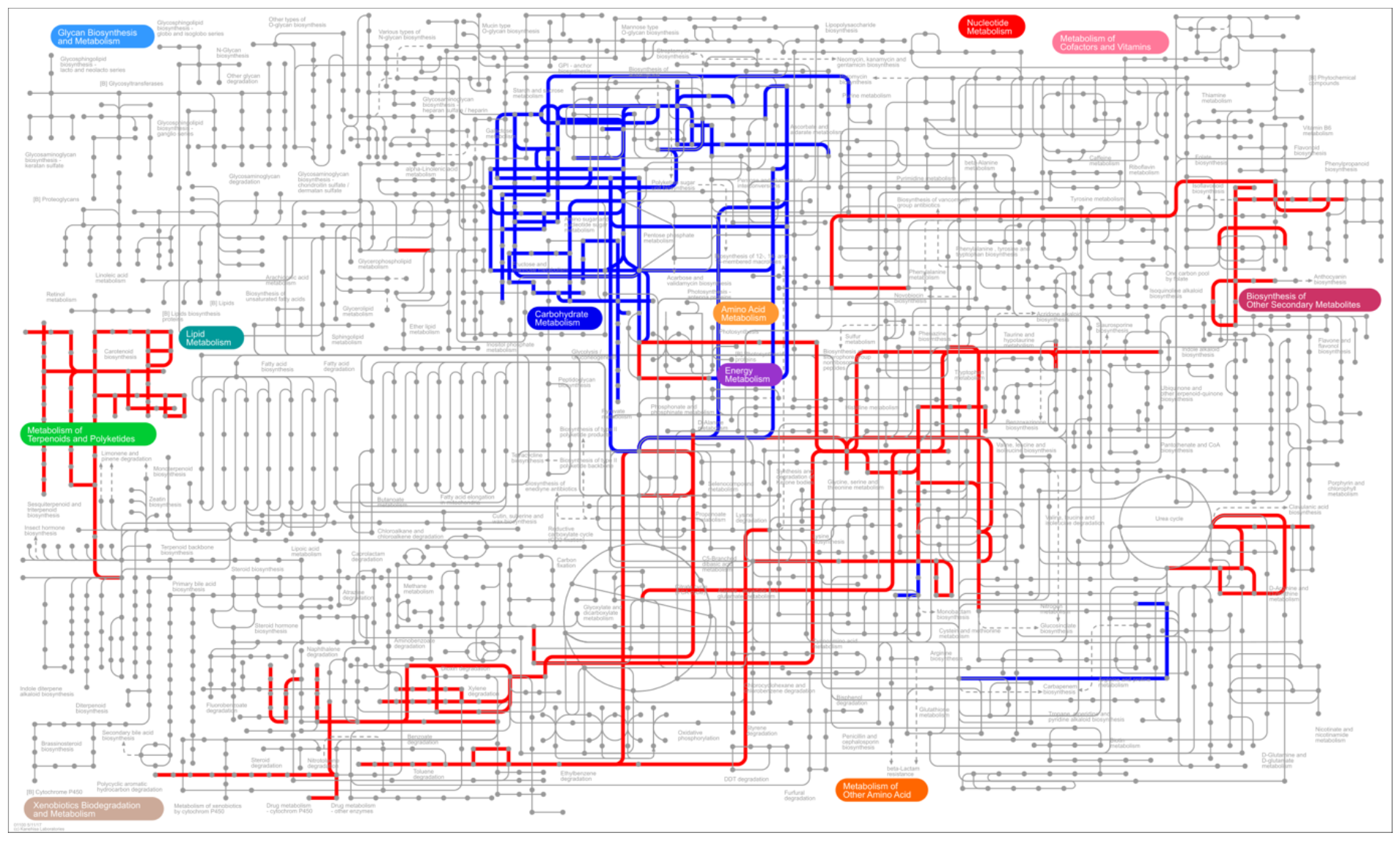
iPath3.0 representation of KEGG metabolic pathways inferred from Tax4Fun2 analysis significantly upregulated in female (red) and male (blue) CRC patients.

